# Machine learning prediction of antimicrobial resistance in respiratory metagenomes

**DOI:** 10.64898/2026.11.20.768127

**Authors:** Michael Aaron Sy, Mattia Prosperi, Panayiotis V. Benos, Simone Marini

**Affiliations:** Department of Epidemiology, University of Florida, Gainesville, FL 32603, USA

## Abstract

**Background:** Targeted therapy for patients with suspected respiratory infections traditionally requires bacterial culture followed by phenotypic antimicrobial susceptibility testing (AST), a process that can take up to 5 days. In acute respiratory distress syndrome (ARDS), delays in identifying the causative pathogen and its resistance profile may be especially harmful because patients are critically ill and often receive broad-spectrum empiric antibiotics. The growing burden of antimicrobial resistance (AMR) further complicates clinical decision-making by reducing effective treatment options and increasing the risk of inadequate initial therapy. Culture-free, AI-driven computational metagenomic strategies have the potential to drastically shorten this window, enabling more rapid targeted therapy prescription. To show the feasibility of such approaches, we develop a proof-of-concept framework based on clinical metagenomics.

**Methods:** We simulated respiratory metagenomes using real-world bacterial genomes with associated antimicrobial susceptibility phenotypes. Each simulated metagenome contained a fixed commensal background and one clinically important respiratory pathogen, *Acinetobacter baumannii*, *Klebsiella pneumoniae*, or *Pseudomonas aeruginosa*, at 5% or 10% bacterial read abundance. Realistic reads were simulated for Oxford Nanopore, PacBio, and Illumina sequencing platforms. Random Forest (RF) classifiers were trained and evaluated using five-fold, strain-grouped cross-validation with fold-specific χ²-ranked 13-mer features. ResFinder and KARGA were included as reference-based metagenomics benchmarks and evaluated using the same five held-out folds across the three sequencing platforms and two pathogen-abundance conditions.

**Results:** RF performance varied substantially across species, antibiotics, and sequencing platforms. The strongest and most consistent results were observed for *K. pneumoniae*, with accuracy reaching approximately 0.96 for ceftazidime, 0.93 for amikacin, and 0.92 for ciprofloxacin. Performance for *P. aeruginosa* was more heterogeneous: long-read models performed well for amikacin, reaching 0.89 with PacBio and 0.88 with Nanopore, whereas Illumina performed substantially better for ciprofloxacin, ceftazidime, and levofloxacin, reaching accuracies of approximately 0.87, 0.77, and 0.70, respectively. Across most organism-antibiotic combinations, RF accuracy was numerically higher than the classification accuracies obtained with KARGA and ResFinder, which generally remained near 0.50. Increasing pathogen abundance from 5% to 10% and increasing the number of selected features from 2,500 to 5,000 generally produced modest improvements in classification accuracy, although the magnitude of improvement varied across organism-antibiotic combinations and sequencing platforms.

**Conclusions:** Genome-wide *k*-mer machine learning can recover phenotype-associated antimicrobial resistance signals from simulated respiratory metagenomes, although performance depends strongly on the organism, antibiotic, and sequencing platform. These results represent a simulation-based proof of concept.

## Introduction

Lower respiratory tract infections are a major global cause of morbidity and mortality, accounting for an estimated 489 million incident cases and 2.49 million deaths worldwide in 2019, and severe respiratory infections such as pneumonia are important precipitants of acute respiratory failure and ARDS (GBD 2019 LRI Collaborators, 2022). In intensive care units, lower respiratory tract infections such as pneumonia represent a substantial source of critical illness; recent severe community-acquired pneumonia guidelines define severe Community Acquired Pneumonia (CAP) pragmatically as CAP requiring ICU admission, and population-based data summarized in a recent review suggest that approximately 40% of patients with CAP require hospitalization and about 5% are admitted to the ICU. Because critically ill patients often require immediate treatment before microbiologic confirmation, empiric antibiotic therapy remains central to pneumonia management, with broad-spectrum coverage recommended when hospital-acquired, ventilator-associated, or drug-resistant pathogens are suspected (Dequin et al., 2026). These infections are further complicated by antimicrobial-resistant (AMR) pathogens. Globally, lower respiratory infections accounted for more than 1.5 million deaths associated with bacterial AMR in 2019, and a recent analysis of bacterial pneumonia hospitalizations found that AMR infections were associated with higher in-hospital mortality (18.5% vs. 10.1%) and longer hospital stays (median 14 vs. 8 days), underscoring their impact on clinical outcomes, healthcare utilization, and available treatment options (Antimicrobial Resistance Collaborators, 2022; Oterino-Moreira et al., 2025). Accurate and rapid characterization of the pathogen’s antimicrobial susceptibility profile is therefore central to clinical decision-making.

Traditional diagnostics rely on culture-based antimicrobial susceptibility testing (AST) to isolate the pathogen and determine its resistance or susceptibility to specific antibiotics. The usual workflow involves primary isolation followed by phenotypic AST methods such as disk diffusion, broth microdilution, or gradient (E-test) procedures (Gajic et al., 2022). Even under optimized laboratory conditions, these steps can require up to ∼62 hours to finalize AST results (Humphries et al., 2021; van Belkum et al., 2020), leading to a total “bed-to-bench” turnaround of roughly 3-5 days from specimen collection to actionable susceptibility information (Dulanto Chiang & Dekker, 2020; Marini, Mora, et al., 2022). During this interval, clinicians are forced to treat empirically, often with broad-spectrum regimens that may be unnecessarily aggressive; while the use of common antibiotics risks to be ineffective and might help selecting resistant bacterial strains (Salam et al., 2023; *The Core Elements of Hospital Antibiotic Stewardship Programs*, n.d.).

Metagenomic next-generation sequencing (mNGS) offers an attractive alternative by sequencing DNA directly from clinical specimens such as sputum or bronchoalveolar lavage fluid (BAL). This approach can, in principle, provide an unbiased view of the microbial community and its resistome, independent of culture conditions and pre-specified panels (Batool & Galloway-Peña, 2023). By leveraging both long-read and short-read sequencing technologies, mNGS can capture extensive genomic diversity, including resistance genes encoded on mobile elements and chromosomal loci, and thus has the potential to inform earlier, more tailored antimicrobial therapy. Using metagenomics could therefore shorten the time to a biologically informed AST profile and uncover resistance patterns that are invisible to conventional or targeted molecular tests (Charalampous et al., 2019; Cheng et al., 2022).

In this study, we introduce an integrated bioinformatics workflow that combines simulated respiratory metagenomic sequencing with machine learning to predict antimicrobial resistance in bacterial respiratory infections. We simulated lung microbial communities containing clinically relevant pathogens across Oxford Nanopore Technologies (ONT), PacBio, and Illumina sequencing platforms and evaluated the effects of pathogen abundance and *k*-mer feature depth on prediction performance. ResFinder and KARGA were included as reference-based metagenomics benchmarks under one standardized ONT condition with 10% pathogen abundance. Because no sufficiently large clinical respiratory metagenomic dataset with matched organism-level antimicrobial susceptibility phenotypes was readily available, the present study was designed as a simulation-based proof of concept rather than a clinical external-validation study.

## Methods

The scripts used to prepare pathogen–commensal genome inputs and simulate respiratory metagenomic sequencing data, together with source-genome identifiers and available simulation seed records, are available at https://github.com/Miaaron/AMR-Respiratory-Metagenomic-Simulation. The underlying reference genomes and simulated sequencing files are not included in the repository due to their large size but can be retrieved from their original sources, BV-BRC and NCBI.

### 2.1 Pathogen and Commensal Species Selection

We first defined candidate bacterial pathogens for acute respiratory distress syndrome (ARDs) and lower respiratory tract infections (LRTI) using a targeted literature review, and then restricted this set based on the availability of genomic and phenotypic data in the Bacterial and Viral Bioinformatics Resource Center (BV-BRC), as follows.

We conducted a targeted literature review of adult lower respiratory tract infection and pneumonia studies that reported species-level microbiology from bronchoalveolar lavage, tracheal aspirates, or sputum. Organisms were prioritized based on repeated mention across six relevant studies of severe lower respiratory tract infection, pneumonia, or ARDS in hospitalized or critically ill populations. The resulting panel included *Streptococcus pneumoniae, Acinetobacter baumannii, Pseudomonas aeruginosa, Klebsiella pneumoniae, Staphylococcus aureus, Haemophilus influenzae, Legionella pneumophila, Escherichia coli, Stenotrophomonas maltophilia, Chlamydia pneumoniae, Mycoplasma pneumoniae, Enterobacter aerogenes, Schewanella algae, Sathylococcus haemolyticus, Staphylococcus epidermidis*, and *Moraxella catarrhalis*. This process resulted in a set of 16 candidate respiratory pathogens that were consistently mentioned across the included studies (listed in **Supplementary Table S1**). We cross-referenced the 16 candidate species against the BV-BRC database to assess data availability. For each candidate species, we evaluated the availability of human respiratory isolates with linked whole-genome sequence data and laboratory-confirmed resistant or susceptible antimicrobial susceptibility phenotypes. Species-antibiotic pairs were retained for analysis if they had at least 50 total isolates and at least 20 isolates in each phenotype class. Under this criterion, three species had sufficient usable data: Acinetobacter baumannii, Klebsiella pneumoniae, and Pseudomonas aeruginosa. To define the commensal species, we started from a publicly available cross-sectional study that profiled the lower-airway microbiome of healthy, never-smoker individuals (NCBI BioProject PRJNA945482; bronchoalveolar lavage, 28 samples). Species-level 16S rRNA profiles from this dataset contained 2,777 unique bacterial species. To derive a tractable yet biologically grounded commensal panel, we cross-referenced these taxa with species reported in independent lower-airway metagenomic or 16S surveys of non-pneumonic lungs and airways (respiratory studies R1-R5 in **Supplementary Table S3**).

To define the commensal background, we used a publicly available dataset of bronchoalveolar lavage samples from healthy, never-smoker individuals (NCBI BioProject PRJNA945482). The dataset included 28 samples and approximately 2,777 identified bacterial taxa. We used this dataset as the initial reference for identifying potential commensal species and considered additional respiratory and oral microbiome studies to identify organisms commonly reported in these communities. From these sources, we selected 87 candidate commensal species for the simulation. However, suitable NCBI reference genomes were available for only 85 species, which were subsequently included in the simulated microbial background. The included species are listed in **Supplementary Table S3**, and their corresponding sequence accession identifiers are available in the GitHub repository.

AMR phenotype data were obtained from the Bacterial and Viral Bioinformatics Resource Center (BV-BRC)(Shukla et al., 2026). We included only bacterial isolates for which (a) the host was “Human” or “Homo sapiens”, (b) annotation confirmed respiratory-related sources, such as the lung, bronchoalveolar lavage (BAL), sputum, tracheal, or bronchial regions, and (c) an associated AST was present from broth microdilution or disk diffusion. After filtering, we retained a total of 184 *A. baumannii*, 2,161 *K. pneumoniae*, and 533 *P. aeruginosa* genomes with paired AST results across the selected organism-antibiotic combinations (per-pathogen and per-antibiotic counts in **Supplementary Table S2**).

### 2.2 Cohort derivation and composition

Following the initial selection and filtering of genomes from BV-BRC, three respiratory pathogens were retained for analysis: *Acinetobacter baumannii*, *Klebsiella pneumoniae*, and *Pseudomonas aeruginosa*. The source dataset contained 184 *A. baumannii*, 2,161 *K. pneumoniae*, and 533 *P. aeruginosa* genome–antibiotic records, corresponding to 184, 419, and 188 unique genome IDs, respectively. The final panel consisted of 11 organism–antibiotic combinations: amikacin for *A. baumannii*; amikacin, cefepime, ceftazidime, ciprofloxacin, levofloxacin, and piperacillin for *K. pneumoniae*; and amikacin, ceftazidime, ciprofloxacin, and levofloxacin for *P. aeruginosa*.

Across the organism–antibiotic combinations retained for modeling, the BV-BRC source dataset contained 1,886 resistant and 992 susceptible genome–antibiotic records. These records were primarily from *K. pneumoniae* (R = 1,553; S = 608), followed by *P. aeruginosa* (R = 187; S = 346) and *A. baumannii* (R = 146; S = 38) (**Supplementary Table S2**). The corresponding per-antibiotic counts were: *A. baumannii*: amikacin (R = 146, S = 38); *K. pneumoniae*: amikacin (R = 42, S = 298), cefepime (R = 248, S = 92), ceftazidime (R = 363, S = 32), ciprofloxacin (R = 347, S = 57), levofloxacin (R = 286, S = 62), and piperacillin (R = 267, S = 67); and *P. aeruginosa*: amikacin (R = 31, S = 147), ceftazidime (R = 52, S = 72), ciprofloxacin (R = 28, S = 32), and levofloxacin (R = 76, S = 95).

Because the available resistant and susceptible isolates were imbalanced for several organisms-antibiotic combinations, class balancing was performed during metagenomic simulation. Within each organism–antibiotic–sequencing platform–pathogen abundance setting, the phenotype with fewer isolates was identified as the minority class. Isolates from this minority class were then used to generate additional independent metagenomic read simulations until the number of simulated samples was approximately equal to that of the majority class. Thus, the same biological isolate could contribute more than one simulated metagenome, with each simulation representing an independent stochastic realization of sequencing reads and community composition. No majority-class isolates were removed, and no synthetic feature-space oversampling methods such as SMOTE were used.

### 2.3 Simulation of Sequencing Reads

To simulate metagenomic sequencing reads, we constructed a lung microbial community consisting of a constant background of 85 commensal bacterial reference genomes and a single pathogen strain introduced at two relative abundances, 5% and 10% of the total microbial community reads. Because clinical respiratory specimens can contain a large excess of host-derived DNA, we modeled the effective microbial fraction after computational or experimental removal of host DNA rather than explicitly simulating human reads. This distinction is particularly relevant for lower-respiratory specimens, in which host-derived reads may comprise the vast majority of shotgun sequencing data; for example, untreated bronchoalveolar lavage samples have been reported to contain a median of approximately 99.7% host reads (Kim et al., 2024). Each simulated microbial community was therefore standardized to an effective non-host sequencing budget of approximately1 x 10^9^ bases (1 Gb). Gigabase-scale sequencing has previously been used as a shallow shotgun metagenomic depth capable of providing informative microbial community profiles (Hillmann et al., 2018), and respiratory mNGS studies have reported sequencing depths ranging from approximately 2 to 25 million reads per sample (Diao et al., 2021). For comparison, 1 Gb corresponds to approximately 6.7 million 150-bp reads, or approximately 3.3 million 150-bp paired-end read pairs, placing the simulated microbial sequencing budget within the scale of sequencing depths used in respiratory metagenomic studies.

The 5% and 10% pathogen abundance conditions were selected as standardized low to moderate pathogen enrichment scenarios rather than as fixed clinical diagnostic thresholds. Respiratory microbial communities associated with pneumonia and acute respiratory distress syndrome (ARDS) are characterized by loss of commensal diversity and enrichment of potentially pathogenic taxa (Kitsios et al., 2020; Montassier et al., 2023). In bronchoalveolar lavage fluid, relative abundances around 5% have also been evaluated as informative thresholds for distinguishing common respiratory pathogens from background organisms, although optimal thresholds vary by organism and clinical setting (Chen et al., 2023). Furthermore, Dickson et al. (2020) observed that Enterobacteriaceae represented approximately 12.5% of bacterial sequences in ARDS specimens compared with 0.18% in non-ARDS specimens, demonstrating that pathogen-associated taxa can occur at abundances on the same order as our higher simulation condition. More severe bacterial pneumonia can progress to pathogen-dominated communities in which a clinically identified pathogen comprises more than 50% of the respiratory bacterial community (Kitsios et al., 2018). Thus, the 5% and 10% conditions were intended to evaluate AMR prediction when the pathogen was enriched but remained a minority component of a complex microbial background. For each simulated community, the total microbial sequencing budget was denoted by *B_m_*= 1 × 10^9^ bases. For a pathogen relative abundance (p), the pathogen-specific sequence allocation was calculated as *B_P_* = *pB_M_*. Consequently, the 5% condition contained approximately 5 × 10 pathogen-derived bases (50 Mb), whereas the 10% condition contained approximately 1 × 10^8^ pathogen-derived bases (100 Mb). The remaining 950 Mb or 900 Mb, respectively, was distributed among the 85 commensal reference genomes. Because sequencing coverage is determined by the number of sequenced bases divided by genome length (Sims et al., 2014), these allocations would correspond, for example, to approximately 10× and 20× coverage for a 5-Mb pathogen genome in the 5% and 10% conditions, respectively; the exact pathogen coverage varied according to genome size. For the commensal community, initial genome-specific allocations were proportional to genome length, randomly perturbed within ±10%, and subsequently rescaled so that the combined commensal allocation equaled the required non-pathogen fraction. Genome-specific coverage was then calculated by dividing the allocated number of bases by the corresponding genome length.

Reads representative of three commonly used sequencing technologies Oxford Nanopore Technologies (ONT), Pacific Biosciences (PacBio), and Illumina were simulated. ONT and PacBio long reads were generated using PBSIM2 (Ono et al., 2021). Genome-specific coverage values were supplied using the --depth parameter. PBSIM2 model-based simulation generates long-read lengths from a modeled distribution and uses hidden Markov models trained on empirical long-read data to reproduce non-uniform quality-score and sequencing-error patterns. Platform-specific HMM model files were supplied using --hmm_model. Simulations used substitution:insertion:deletion difference ratios were 23:31:46 for ONT and 6:50:54 for PacBio. PBSIM2 defines --depth as target genome coverage and --difference-ratio in substitution:insertion:deletion order.

Illumina reads were simulated using InSilicoSeq (Gourlé et al., 2019) with its built-in NovaSeq error model, producing 150-bp paired-end reads in separate R1 and R2 FASTQ files. To facilitate comparisons across sequencing technologies, the genome-specific coverage profile generated for each ONT pathogen–commensal community was used as the reference coverage profile for the corresponding Illumina and PacBio simulations. For Illumina, these coverage values were supplied to InSilicoSeq using the --coverage_file option, which permits explicit specification of genome-level sequencing coverage. The same genome-specific coverage targets were used for the corresponding PacBio simulations. Consequently, ONT, PacBio, and Illumina simulations represented approximately equivalent pathogen and commensal genome-level coverage while retaining platform-specific read lengths and sequencing-error characteristics.

### 2.4 Analytical Datasets

The primary analysis included 11 organism-antibiotic combinations representing three respiratory pathogens. The analytical panel consisted of *A. baumannii* (amikacin); *K. pneumoniae* (amikacin, cefepime, ceftazidime, ciprofloxacin, levofloxacin, and piperacillin); and *P. aeruginosa* (amikacin, ceftazidime, ciprofloxacin, and levofloxacin). Each organism-antibiotic dataset included simulated metagenomes derived from isolates with resistant or susceptible phenotypes obtained from BV-BRC.

Because the numbers of resistant and susceptible isolates varied substantially across organism antibiotic combinations, the simulation procedure was also used to reduce phenotype class imbalance. Within each organism antibiotic, sequencing platform, and pathogen abundance setting, additional metagenomes were generated from isolates belonging to the underrepresented phenotype until the numbers of resistant and susceptible simulated samples were approximately balanced. Balancing was therefore performed through additional independent simulations rather than by discarding isolates, relabeling samples, or applying synthetic feature-space oversampling.

Replicate simulations were generated using independent simulator random seeds while retaining the phenotype and underlying pathogen genome of the source isolate. To prevent replicate-level information leakage during model evaluation, all simulations originating from the same isolate were assigned to the same cross-validation fold.

For the primary Random Forest (RF) experiment, each eligible isolate was represented across three sequencing platforms-ONT, PacBio, and Illumina-and two pathogen-abundance levels, 5% and 10%. Models were evaluated using the top 2,500 and 5,000 fold-specific χ²-ranked 13-mer features. This produced 12 primary simulation settings per organism-antibiotic combination: three sequencing platforms, two abundance levels, and two feature-selection depths. Across 11 organism-antibiotic combinations, the complete factorial analysis therefore comprised 132 RF model settings, each evaluated using five strain-grouped held-out folds.

ResFinder and KARGA were evaluated as reference-based metagenomics benchmarks across the three sequencing platforms (ONT, PacBio, and Illumina) and at both 5% and 10% pathogen abundance. Predictions from both methods were evaluated using the same five strain-grouped held-out test partitions used for the corresponding RF models. Since these reference-based methods do not require model training, the fold structure was used to evaluate their performance on matched test sets rather than for cross-validation during model fitting.

The organism-antibiotic combinations and their corresponding isolate and simulation characteristics are summarized in **Supplementary Table S2**. The genome preparation and simulation scripts, source-genome identifiers, and available simulation seed records are provided in the GitHub repository at https://github.com/Miaaron/AMR-Respiratory-Metagenomic-Simulation.

### 2.5 Sequence Data Processing

Illumina sequencing outputs (R1 and R2) were concatenated at the file level to combine paired-end reads (one file per sample). For ONT and PacBio sequencing, the outputs were merged into a single consolidated FASTQ file (one per sample). These unified FASTQ files formed the input for downstream feature extraction and AMR prediction.

### 2.6 *k*-mer Feature Matrix Creation and Selection

To identify genomic patterns associated with antimicrobial resistance, we represented each simulated metagenomic sample using *k*-mers. We define *k*-mers as all contiguous nucleotide subsequences of length k extracted from a DNA sequence. Each sequencing sample was represented by the presence or abundance of these fixed-length sequence patterns, which can capture resistance-associated genes, mutations, and broader genomic backgrounds without requiring prior alignment to a reference genome. Of note, *k*-mers are used in Bioinformatics applied to AMR prediction, but notably, existing algorithms are designed for cultured samples or environmental metagenomics (Bortolaia et al., 2020; Marini et al., 2023; Marini, Oliva, et al., 2022; Prosperi & Marini, 2021; Serajian et al., 2024). We selected k=13 based on our previous work(Marini, Oliva, et al., 2022). We extracted pathogen-specific *k*-mer databases directly from the BV-BRC genomes for *A. baumannii*, *K. pneumoniae*, and *P. aeruginosa* using KMC3 (Kokot et al., 2017). For each pathogen, all unique 13-mers present in the corresponding reference genomes were counted, resulting in pathogen-specific *k*-mer sets of ∼10-15 million *k*-mers. We performed univariate feature selection using chi-square (χ²) tests for *k*-mer feature selection. Feature selection was always performed only on the training data within each fold, and the selected *k*-mers were then applied to the held-out test data, thereby preventing information leakage from the test set into the feature selection process. For each pathogen-antibiotic combination, feature selection was performed separately within each training fold using chi-square tests. The top 2,500 and 5,000 ranked 13-mers were then evaluated as separate feature sets using the corresponding held-out test data.

### 2.7 Machine Learning Model and Cross-Validation

A RF classifier was trained to predict AMR status based on the selected *k*-mer features. For each of the 11 pathogen-antibiotic models, we first assembled a dataset that included all simulated metagenomic samples for that specific combination, all resistant and susceptible simulations generated from the BV-BRC genomes at 5% and 10% pathogen abundance (per-pathogen and per-antibiotic sample counts are reported in **Supplementary Table S2**). Because the number of available resistant and susceptible genomes differed across pathogen-antibiotic pairs, the total number of samples per model varied accordingly.

For comparison with current reference-based metagenomics AMR detection tools, we evaluated ResFinder version 2.4.0 and KARGA. Both benchmark methods were evaluated under a standardized condition consisting of ONT simulated reads at 10% pathogen abundance and were applied to the same strain-grouped held-out test partitions used for the corresponding RF models. Performance was calculated independently within each fold and summarized across the five folds, allowing direct comparison of the methods on identical test samples. Because ResFinder and KARGA do not require statistical model training, the fold structure was used to provide matched test-set evaluation rather than cross-validation for model fitting.

For comparison with reference-based metagenomics AMR detection tools, we evaluated ResFinder version 2.4.0 and KARGA. Both methods were applied to simulated reads generated using ONT, PacBio, and Illumina at 5% and 10% pathogen abundance. Their predictions were evaluated using the same strain-grouped held-out test partitions as the corresponding RF models. Performance was calculated separately for each fold and summarized across the five folds.

## Results

### Primary analysis: *k*-mer machine learning versus reference-based-based methods

Across all three sequencing platforms, the RF models generally outperformed the reference-based-based approaches, particularly for *Klebsiella pneumoniae* and *Acinetobacter baumannii*. Increasing the number of selected k-mers from 2,500 to 5,000 generally produced only modest changes in performance, and the effect of increasing pathogen abundance from 5% to 10% varied by organism and antibiotic. In contrast, KARGA and ResFinder performance was typically concentrated near 0.50, with relatively little improvement at higher pathogen abundance.

#### Oxford Nanopore Technologies

For ONT sequencing, RF performance was strongest for *K. pneumoniae* and remained comparatively lower for several *Pseudomonas aeruginosa* antibiotics (Figure 1). At 5% pathogen abundance, RF accuracy using 5,000 selected features ranged from 0.79 for *K. pneumoniae* cefepime to 0.95 for ceftazidime. At 10% abundance, performance remained similar, ranging from 0.78 for cefepime to 0.95 for ceftazidime; amikacin and ciprofloxacin achieved accuracies of 0.93 and 0.91, respectively. For *A. baumannii* amikacin, accuracy was 0.80 at 5% abundance and 0.76 at 10% abundance using 5,000 features.

**Figure 1.**
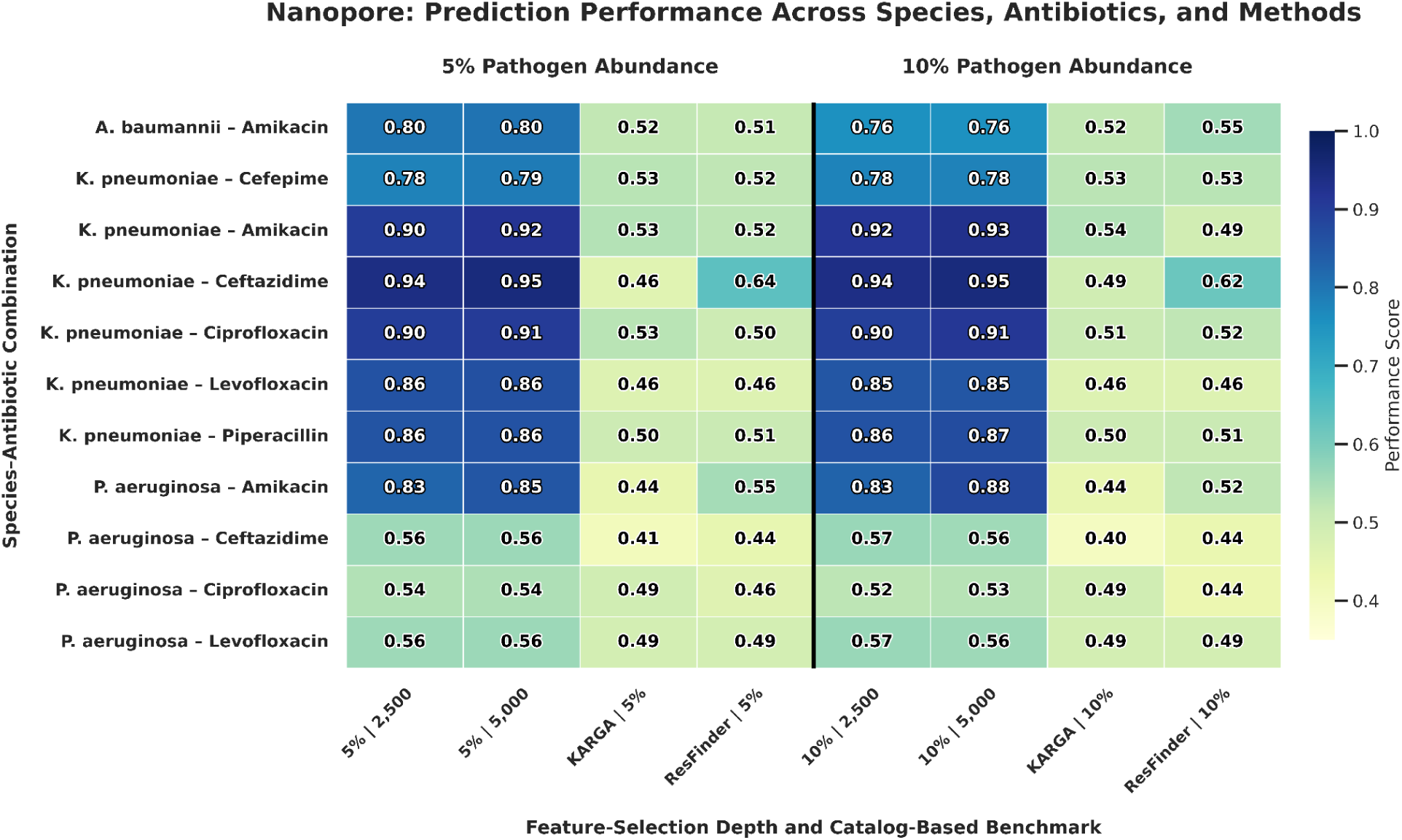
ONT prediction performance across species, antibiotics, pathogen abundance, feature depth, and reference methods. RF models were evaluated at 5% and 10% pathogen abundance using 2,500 and 5,000 fold-specific χ²-ranked 13-mers. KARGA and ResFinder were evaluated at both pathogen-abundance levels. RF values represent accuracy, whereas KARGA and ResFinder values represent classification accuracy from the corresponding reference-based-based benchmark outputs.

Performance for *P. aeruginosa* was more heterogeneous. At the 10% abundance and 5,000-feature setting, accuracy reached 0.88 for amikacin but was substantially lower for ceftazidime, ciprofloxacin, and levofloxacin at 0.56, 0.53, and 0.56, respectively. Similar patterns were observed at 5% abundance.

KARGA and ResFinder generally performed close to chance-level classification for ONT data. At 5% abundance, KARGA values ranged from 0.41 to 0.53 and ResFinder values from 0.44 to 0.64. At 10% abundance, KARGA ranged from 0.40 to 0.54 and ResFinder from 0.44 to 0.62. The strongest reference-based-based result was observed for ResFinder in *K. pneumoniae* ceftazidime, with performance of 0.64 at 5% and 0.62 at 10% abundance.

#### PacBio

PacBio produced the strongest RF performance overall for several organism-antibiotic combinations (Figure 2). For *A. baumannii* amikacin, accuracy increased from 0.84 at 5% abundance to 0.85 at 10% abundance using 5,000 features. RF performance for *K. pneumoniae* was consistently high. At 10% abundance and 5,000 features, accuracy was 0.82 for cefepime, 0.93 for amikacin, 0.96 for ceftazidime, 0.92 for ciprofloxacin, 0.88 for levofloxacin, and 0.87 for piperacillin. Similar values were observed at 5% pathogen abundance.

**Figure 2.**
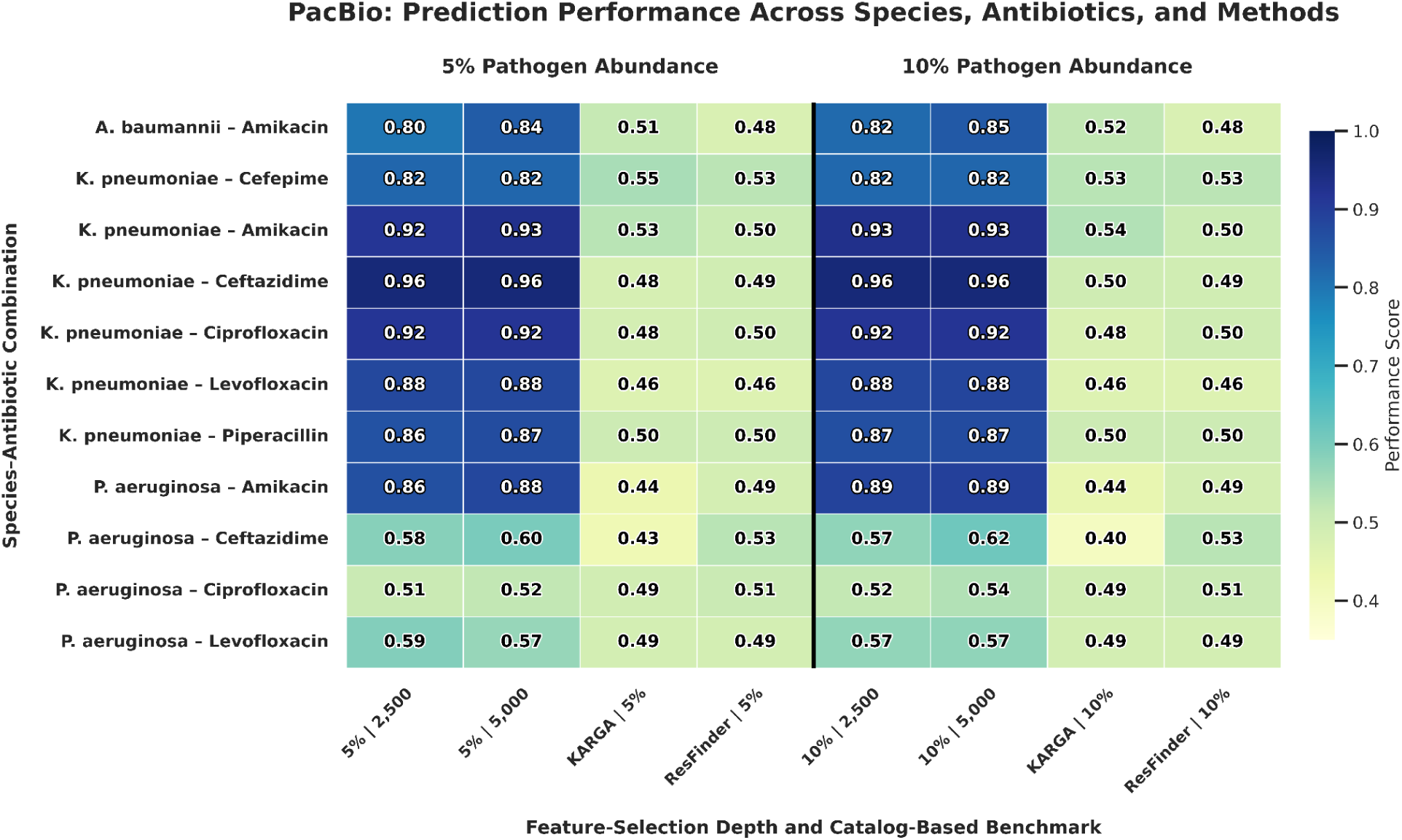
PacBio prediction performance across species, antibiotics, pathogen abundance, feature depth, and reference methods. RF models were evaluated at 5% and 10% pathogen abundance using 2,500 and 5,000 fold-specific χ²-ranked 13-mers. KARGA and ResFinder were evaluated at both pathogen-abundance levels. RF values represent accuracy, whereas KARGA and ResFinder values represent classification accuracy from the corresponding reference-based-based benchmark outputs.

For *P. aeruginosa*, PacBio RF performance varied substantially by antibiotic. Amikacin achieved accuracy of 0.88 at 5% abundance and 0.89 at 10% abundance using 5,000 features. In contrast, ceftazidime, ciprofloxacin, and levofloxacin reached 0.62, 0.54, and 0.57, respectively, at the 10% abundance and 5,000-feature setting.

reference-based-based performance was again substantially lower than RF performance for most combinations. At 5% abundance, KARGA ranged from 0.43 to 0.55 and ResFinder from 0.46 to 0.53. At 10% abundance, KARGA ranged from 0.40 to 0.54 and ResFinder from 0.46 to 0.53. Neither increased abundance nor organism-specific differences produced a consistent improvement in reference-based-based performance.

#### Illumina

Illumina RF performance was more variable for *K. pneumoniae* than PacBio or ONT but was notably strong for several *P. aeruginosa* antibiotics (Figure 3). For *A. baumannii* amikacin, accuracy was 0.76 at 5% abundance and 0.73 at 10% abundance using 5,000 features. Among *K. pneumoniae* combinations at 10% abundance and 5,000 features, accuracy ranged from 0.58 for cefepime to 0.86 for ceftazidime. Corresponding values were 0.79 for amikacin, 0.77 for ciprofloxacin, 0.77 for levofloxacin, and 0.72 for piperacillin.

**Figure 3.**
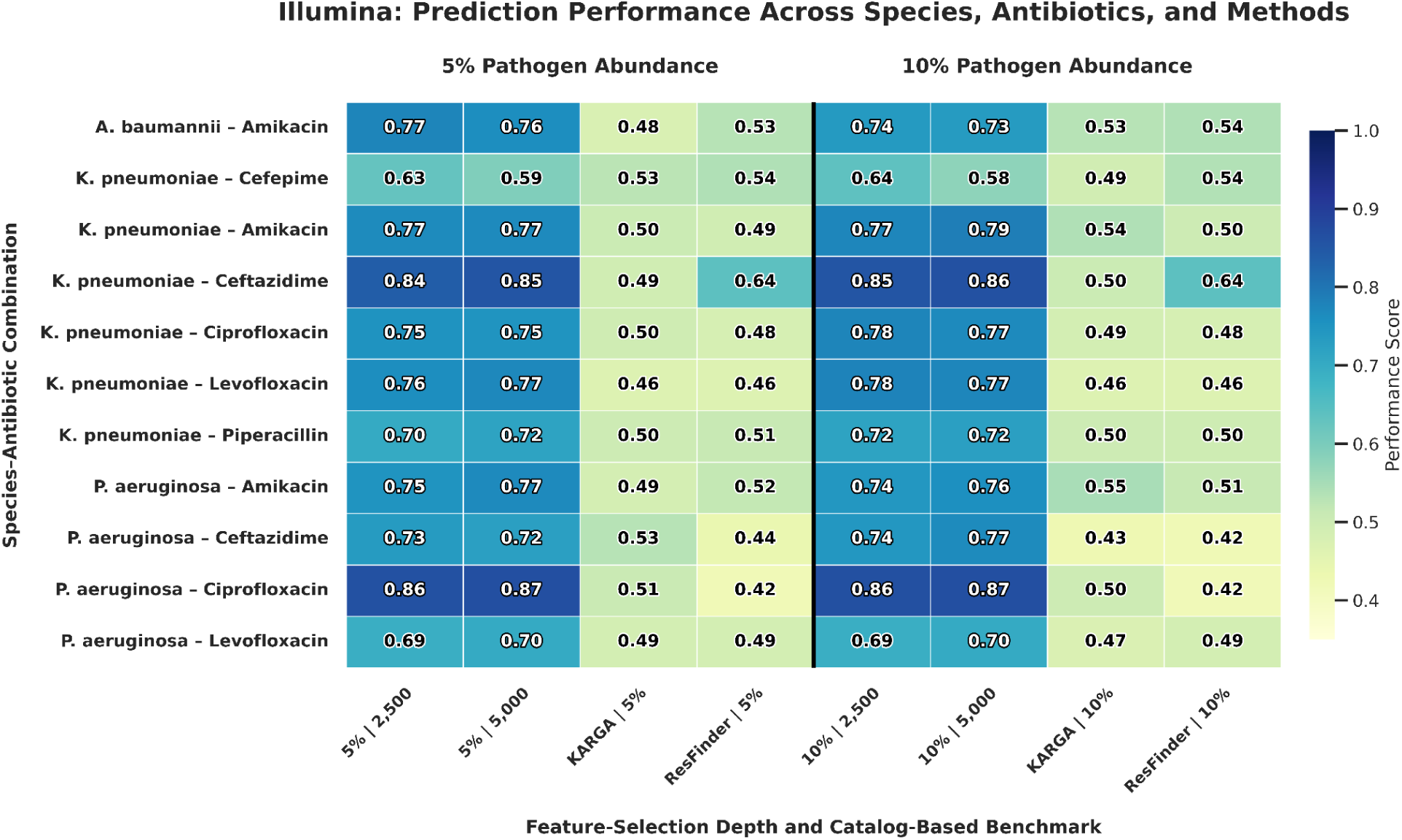
Illumina prediction performance across species, antibiotics, pathogen abundance, feature depth, and reference methods. RF models were evaluated at 5% and 10% pathogen abundance using 2,500 and 5,000 fold-specific χ²-ranked 13-mers. KARGA and ResFinder were evaluated at both pathogen-abundance levels. RF values represent accuracy, whereas KARGA and ResFinder values represent classification accuracy from the corresponding reference-based-based benchmark outputs.

Illumina performed particularly well for *P. aeruginosa* ciprofloxacin. At both 5% and 10% pathogen abundance, the 5,000-feature model achieved accuracy of 0.87, substantially exceeding the corresponding ONT and PacBio values. At 10% abundance, accuracy was also 0.76 for amikacin, 0.77 for ceftazidime, and 0.70 for levofloxacin.

KARGA and ResFinder again showed considerably lower performance than RF. At 5% abundance, KARGA values ranged from 0.46 to 0.53 and ResFinder from 0.42 to 0.64. At 10% abundance, KARGA ranged from 0.43 to 0.55 and ResFinder from 0.42 to 0.64. As observed for ONT, ResFinder achieved its strongest result for *K. pneumoniae* ceftazidime, with an accuracy of 0.64 at both abundance levels.

#### Overall comparison across platforms and methods

Across the three sequencing technologies, RF performance depended more strongly on the organism-antibiotic combination than on either pathogen abundance or feature-selection depth. PacBio produced the strongest results for most *K. pneumoniae* combinations, including a maximum accuracy of 0.96 for ceftazidime. ONT also performed strongly for *K. pneumoniae*, whereas Illumina showed the strongest performance for *P. aeruginosa* ciprofloxacin, reaching 0.87 at both pathogen-abundance levels with 5,000 selected features.

In contrast, KARGA and ResFinder generally produced performance values near 0.50 across platforms, abundance levels, and antibiotics. The limited change between 5% and 10% abundance suggests that increasing pathogen representation alone did not substantially improve reference-based-based resistance classification in these simulated metagenomes. Overall, the *k*-mer-based RF models provided substantially greater discrimination than the reference-based-based approaches for most organism-antibiotic combinations, although performance remained challenging for several *P. aeruginosa* phenotypes.

### Effect of feature depth and pathogen abundance

Increasing the number of selected *k*-mers from 2,500 to 5,000 produced some improvements, but the magnitude of improvement was generally modest and not consistent across all settings. For example, at 10% pathogen abundance, *P. aeruginosa* amikacin using ONT improved from 0.83 to 0.88, and *P. aeruginosa* ceftazidime using PacBio improved from 0.57 to 0.62. However, many other combinations showed little change or even slight decreases. For example, *K. pneumoniae* cefepime using Illumina decreased from 0.64 to 0.58 when increasing from 2,500 to 5,000 features.

Similarly, increasing pathogen abundance from 5% to 10% did not uniformly improve performance. Some combinations improved slightly, while others remained stable or decreased. For instance, *P. aeruginosa* amikacin using ONT increased from 0.85 to 0.88 at 5,000 features, but *A. baumannii* amikacin using ONT decreased from 0.80 to 0.76. This suggests that higher pathogen abundance alone did not guarantee better model performance. Instead, performance appears to depend on the interaction between species, antibiotic, sequencing platform, and the *k*-mer feature set.

### Platform- and antibiotic-specific patterns

Overall, PacBio often produced the highest or near-highest RF performance, especially for *K. pneumoniae* and for *P. aeruginosa* amikacin. ONT also performed well for many *K. pneumoniae* antibiotics and for *P. aeruginosa* amikacin, indicating that long-read sequencing can support strong *k*-mer-based resistance prediction for certain organism-drug combinations.

Illumina showed a more mixed pattern. It performed lower than ONT and PacBio for several *K. pneumoniae* antibiotics and for *A. baumannii* amikacin, but it was the best-performing platform for *P. aeruginosa* ciprofloxacin and levofloxacin. This suggests that short-read sequencing may better capture certain resistance-associated signals for *P. aeruginosa* fluoroquinolone resistance under these simulation conditions.

Taken together, these results support χ²-selected 13-mer RF models as a strong alternative to ResFinder-based reference-based metagenomics prediction. However, the results also show that performance is not driven by a single factor such as pathogen abundance, feature depth, or sequencing platform alone. Instead, prediction accuracy depends on the combined effects of organism, antibiotic, platform, and *k*-mer representation.

### Error-free reads

To determine whether the comparatively low performance of the long-read models for *Pseudomonas aeruginosa* was primarily attributable to sequencing errors, we conducted an additional experiment using error-free, or “perfect,” simulated ONT reads. All baseline simulation parameters were retained from the corresponding primary ONT experiments, including the pathogen and commensal genomes, community composition, pathogen abundance, genome-specific coverage targets, and downstream analytical pipeline. The only intentional modification was the removal of simulator-introduced substitutions, insertions, and deletions. Thus, the experiment isolated the contribution of sequencing error while holding the underlying genomic composition and sequencing-depth assumptions constant. RF models were evaluated using the same *k*-mer extraction, χ² feature-selection, strain-grouped cross-validation, and Random Forest procedures as the primary analysis, with the top 2,500 and 5,000 ranked 13-mer features evaluated separately.

Removing sequencing errors did not substantially improve prediction. For ceftazidime, mean accuracy was 0.53 using both 2,500 and 5,000 features. For ciprofloxacin, accuracy was 0.47 and 0.50, respectively, while for levofloxacin it was 0.55 and 0.56. These results were similar to those obtained using the original ONT simulations, in which accuracy remained in the range of 0.52-0.57 for the same species-antibiotic combinations.

Therefore, simulator-introduced sequencing error was not the primary explanation for the weak long-read performance observed for these *P. aeruginosa* phenotypes. Instead, the limiting factors may include insufficient coverage of phenotype-informative genomic regions, inadequate representation of resistance-associated variants among the selected *k*-mers, limitations of univariate feature selection, or antibiotic-specific resistance mechanisms that are not fully captured by untargeted *k*-mer presence. The comparatively strong long-read performance for *P. aeruginosa* amikacin further indicates that the reduced performance was antibiotic-specific rather than a general inability to model resistance in *P. aeruginosa*.

## Discussion

We performed a simulation-based study on real-world genomes to investigate whether traditional reference-based approaches and untargeted *k*-mer-based machine learning methods can accurately predict antimicrobial resistance (AMR) phenotypes from clinical metagenomic sequencing data. Our main finding is that the traditional metagenomics AMR prediction approaches (KARGA, ResFinder) do seem to provide an acceptable performance when used to infer AMR in clinical metagenomics samples. On the other hand, untargeted *k*-mer-based machine learning can recover antimicrobial resistance associated genomic signals that are not consistently captured by current reference-based metagenomics based approaches. However, its predictive performance varied markedly across pathogens, antibiotic combinations and sequencing platforms. RF models trained on χ²-ranked 13-mer features performed particularly well for *Klebsiella pneumoniae*, with accuracies ranging from 0.82 for cefepime to 0.96 for ceftazidime. These findings indicate that genome wide *k*-mer profiles contain substantial phenotype-associated information for several clinically important *K. pneumoniae* resistance phenotypes and support the potential value of data driven genomic features as a complement to predefined resistance gene reference-baseds.

Predictive performance was also comparatively strong for *A. baumannii* amikacin resistance, with the highest accuracy reaching approximately 0.85 for PacBio, compared with approximately 0.80 for ONT and 0.77 for Illumina. Taken together with the *K. pneumoniae* results, these findings suggest that resistance associated genomic signals can remain detectable even when pathogen reads are embedded within a simulated polymicrobial background rather than analyzed as isolated pathogen genomes. This is important for metagenomic AMR prediction, where clinically relevant resistance determinants must be identified in the presence of a substantial nonpathogen sequence.

Another important finding of our study was that no sequencing platform was uniformly superior across all organism × antibiotic combinations. Relative performance depended on the specific resistance phenotype being modeled: long reads performed particularly well for several *K. pneumoniae* phenotypes and for *P. aeruginosa* amikacin resistance, whereas Illumina performed better for several other *P. aeruginosa* phenotypes. These differences suggest that platform performance is shaped not only by sequencing technology itself, but also by the genomic architecture of resistance, the abundance and distribution of phenotype informative sequence features, and the extent to which those features are preserved under platform specific read characteristics. The weaker performance observed for several *P. aeruginosa* phenotypes therefore appears to reflect organism and antibiotic-specific limitations rather than a general failure of the *k*-mer framework or of any single sequencing platform.

The comparatively inferior and more variable performance observed for several *P. aeruginosa* phenotypes warrants separate consideration, because these results differed from the stronger and more consistent patterns seen for *K. pneumoniae* and *A. baumannii*. One possible explanation is that resistance to ciprofloxacin and levofloxacin may depend heavily on a limited number of specific chromosomal variants. If reads do not cover the relevant loci, or if the corresponding *k*-mers are not retained during fold-specific feature selection, the predictive signal may be lost even when the reads themselves are error-free.

Ceftazidime resistance may present a different challenge because it can reflect combinations of acquired resistance genes, chromosomal mutations, porin alterations, efflux activity, and regulatory changes. Some mechanisms may be difficult to distinguish using simple *k*-mer presence or absence, particularly when the relevant genes are present in both susceptible and resistant isolates but differ in regulation or expression.

The error-free ONT experiment was designed to test whether sequencing errors disrupted resistance-associated *k*-mers and thereby reduced classification performance. If sequencing error had been the primary limiting factor, removing those errors would have been expected to produce a substantial increase in accuracy. Instead, performance remained approximately 0.47-0.56. This finding indicates that base-level sequencing noise alone does not explain the poor long-read performance.

The stronger Illumina results suggest that the relevant genomic signal was present in the source genomes but was represented more effectively in the short-read feature matrices. However, the error-free experiment indicates that this advantage cannot be explained solely by lower sequencing error. Differences in effective genome coverage, read sampling, feature sparsity, read length, and the number of observations contributing to each *k*-mer may also influence model performance.

### Comparison with reference-based-based resistance detection

Under the standardized ONT 10% pathogen abundance condition, the *k*-mer RF models generally demonstrated stronger performance than ResFinder and KARGA. All three approaches were evaluated using the same five strain grouped held out folds, providing a fold matched comparison on identical test samples. The observed differences therefore cannot be attributed simply to differences in evaluation partitions.

These findings also highlight important limitations of current existing reference-based metagenomics detection based tools when they are applied directly to clinical metagenomic data. Such approaches depend on the detection of previously characterized resistance genes or variants represented in reference databases and may therefore fail to capture resistance arising from novel mutations, combinations of chromosomal changes, regulatory mechanisms, or broader genomic backgrounds associated with phenotype. Their performance may also be affected when pathogen derived reads constitute only a minority of a polymicrobial sample. Thus, although reference-based metagenomics based tools are valuable for identifying known resistance determinants, they may be insufficient as stand alone methods for comprehensive phenotypic AMR prediction from clinical metagenomes.

Existing reference-based metagenomics detection tools and machine learning approaches should therefore be considered complementary rather than interchangeable. reference-based metagenomics detection methods provide interpretable evidence for recognized resistance determinants, whereas *k*-mer models can potentially capture additional sequence patterns associated with resistance phenotypes. However, the present study used simulated metagenomic sequencing data, and the ability of these models to predict AMR directly from clinical metagenomic specimens remains to be validated.

The present findings should not be interpreted as demonstrating superiority over culture based antimicrobial susceptibility testing. Culture based testing provided the phenotype labels used as the reference outcome and was not evaluated as a competing prediction model. Rather, the potential clinical value of the *k*-mer approach lies in providing earlier resistance prediction directly from sequencing data while conventional phenotypic susceptibility testing is still being completed.

Overall, differences associated with species, antibiotic, and sequencing platforms were larger than the average effects of either feature depth or pathogen abundance. These findings suggest that simply increasing the number of *k*-mers or the simulated pathogen fraction is unlikely to resolve poorly performing combinations. Antibiotic-specific feature engineering and model selection may be more effective.

### Limitations

This study has several limitations. First, the analyses were based on simulated metagenomic communities. Although simulation enables controlled comparisons across sequencing platforms, pathogen-abundance levels, and feature-selection settings, it cannot fully reproduce the biological and technical complexity of clinical respiratory specimens. Factors such as host DNA contamination, variable extraction efficiency, inhibitory substances, uneven genome coverage, mixed-strain infections, and sequencing-batch effects were not represented.

Second, the available phenotype distributions limited the number of species-antibiotic combinations that could be evaluated. The findings may therefore not generalize to other pathogens, antimicrobial agents, resistance mechanisms, or clinical populations.

Third, feature selection was based on univariate χ² testing. This method evaluates each *k*-mer separately and may fail to retain combinations of individually weak but jointly informative markers. It may also select large numbers of correlated *k*-mers originating from the same genomic region. Consequently, the top 2,500 or 5,000 features may not represent the optimal feature set for every phenotype.

Fourth, RF was the primary machine-learning method in the main comparison. Other classifiers, including regularized logistic regression, linear support-vector machines, gradient-boosting approaches, and models designed specifically for sparse genomic features, may produce different results. Antibiotic-specific model selection may be particularly important for *P. aeruginosa*.

Fifth, the error-free analysis removed sequencing errors but did not independently isolate the effects of read length, genome coverage, pathogen-read count, background-community composition, or *k*-mer-generation procedures. The analysis therefore demonstrates that sequencing error was not the primary limiting factor but does not identify the exact source of the remaining performance limitations.

Finally, independent external validation was not completed because sufficiently large clinical respiratory metagenomic datasets containing matched organism-level antimicrobial susceptibility phenotypes were not available. Available clinical microbiome datasets did not provide enough eligible samples for a reliable evaluation of model generalizability. The present results should therefore be interpreted as simulation-based evidence, and performance in clinical specimens remains to be established.

## Conclusion

Genome-wide χ²-ranked *k*-mer features combined with RF classification produced strong antimicrobial resistance predictions for several clinically important species-antibiotic combinations. Performance was particularly high for *K. pneumoniae* especially against amikacin, ciprofloxacin, and levofloxacin, for which conventional resistance gene-based methods (KARGA and ResFinder) performed poorly, with scores generally near 0.50. However, performance varied substantially by species, antibiotic, and sequencing platform.

Overall, these findings demonstrate the potential of genome-wide k-mer models to complement reference-based-based resistance detection while identifying important species-, antibiotic-, and platform-specific limitations. Because sufficiently large clinical metagenomic datasets with matched susceptibility phenotypes were not available for reliable external validation, the results should be considered a simulation-based proof of concept. Additional methodological development and evaluation in larger clinical respiratory cohorts will be necessary before this approach can be considered for antimicrobial decision support.

