## Supplementary Tables S1 - S3 for "Machine learning prediction of antimicrobial resistance in respiratory metagenomes"

**Supplementary Material**

**Supplementary Table S1. Candidate respiratory pathogens and BV-BRC screening observations**

Sixteen candidate organisms were compiled from a targeted review of respiratory infection literature. Historical BV-BRC data-collection notes from October 2023 document the availability and limitations of genome records and resistance phenotypes under several sets of human-host, respiratory-source and laboratory-method filters. The observations below refer to those searches; they do not indicate that a species lacks AMR data across all BV-BRC records.

| Candidate organism | Used in simulations | Historical BV-BRC screening observation |
|---|---|---|
| Streptococcus pneumoniae | No | No genome records returned under the documented BAL-focused filter. |
| Acinetobacter baumannii | Yes | Eligible source genomes and resistance phenotypes were retained. |
| Pseudomonas aeruginosa | Yes | Retained; a restrictive respiratory search had qualifying laboratory-method entries. |
| Klebsiella pneumoniae | Yes | Retained; narrow respiratory searches yielded phenotype data. |
| Staphylococcus aureus | No | No qualifying laboratory-method records in the documented restricted search. |
| Haemophilus influenzae | No | No human-specific AMR records under the documented filters. |
| Legionella pneumophila | No | No AMR phenotypes in the documented search. |
| Escherichia coli | No | Few phenotype records under restrictive respiratory/BAL searches (four in one search). |
| Stenotrophomonas maltophilia | No | Limited usable data; dependent records noted in screening documentation. |
| Chlamydia pneumoniae | No | No AMR phenotypes in the documented search. |
| Mycoplasma pneumoniae | No | No AMR phenotypes in the documented search. |
| Enterobacter aerogenes | No | Limited usable data; dependent records noted in screening documentation. |
| Shewanella algae | No | Limited usable data; dependent records noted in screening documentation. |
| Staphylococcus haemolyticus | No | Limited usable data; dependent records noted in screening documentation. |
| Staphylococcus epidermidis | No | Limited usable data; dependent records noted in screening documentation. |
| Moraxella catarrhalis | No | No AMR phenotypes in the documented search. |

Note: Pathogen screening was performed using BV-BRC in October 2023. Availability of genome sequences and antimicrobial resistance phenotype data was assessed using human-host, respiratory-source, and laboratory-testing filters. Screening results varied depending on the criteria applied. These observations reflect the records available under the documented searches and do not establish the absence of data in BV-BRC more generally.

**Supplementary Table S2. BV-BRC source-genome records by pathogen, antibiotic and phenotype**

Counts were derived from the source-genome manifest used to document genome retrieval for the simulation pipeline. Each genome–antibiotic–phenotype record is counted once for its combination. Individual genome IDs may appear for multiple antibiotics; these totals are not independent-isolate counts or numbers of simulated metagenomes.

| Pathogen | Antibiotic | Resistant | Susceptible | Total |
|---|---|---|---|---|
| Acinetobacter baumannii | amikacin | 146 | 38 | 184 |
| Klebsiella pneumoniae | amikacin | 42 | 298 | 340 |
| Klebsiella pneumoniae | cefepime | 248 | 92 | 340 |
| Klebsiella pneumoniae | ceftazidime | 363 | 32 | 395 |
| Klebsiella pneumoniae | ciprofloxacin | 347 | 57 | 404 |
| Klebsiella pneumoniae | levofloxacin | 286 | 62 | 348 |
| Klebsiella pneumoniae | piperacillin | 267 | 67 | 334 |
| Pseudomonas aeruginosa | amikacin | 31 | 147 | 178 |
| Pseudomonas aeruginosa | ceftazidime | 52 | 72 | 124 |
| Pseudomonas aeruginosa | ciprofloxacin | 28 | 32 | 60 |
| Pseudomonas aeruginosa | levofloxacin | 76 | 95 | 171 |
| Total | | 1886 | 992 | 2878 |

Distinct BV-BRC genome IDs represented: Acinetobacter baumannii, 184; Klebsiella pneumoniae, 419; Pseudomonas aeruginosa, 188 (total 791 distinct IDs). The complete per-genome listing and phenotypes are available in the repository: https://github.com/Miaaron/AMR-Respiratory-Metagenomic-Simulation.

**Supplementary Table S3. Commensal bacterial reference-genome species used in simulation**

The commensal background was constructed using a pragmatic selection of species from a healthy-participant BAL microbiome dataset (NCBI BioProject PRJNA945482) and a small number of related respiratory and oral microbiome studies. Eighty-seven candidate species were initially considered; according to the study records, suitable NCBI reference genomes were available for 85, which were included in the simulations. The 85 species are listed below; recovered NCBI sequence accessions are available in the GitHub repository.

| Commensal species |
|---|
| Aggregatibacter segnis |
| Alloprevotella rava |

| **Commensal species** |
| --- |
| Alloprevotella tannerae |
| Arachnia propionica |
| Bifidobacterium dentium |
| Campylobacter concisus |
| Campylobacter rectus |
| Capnocytophaga gingivalis |
| Capnocytophaga granulosa |
| Capnocytophaga ochracea |
| Capnocytophaga sputigena |
| Cardiobacterium hominis |
| Corynebacterium accolens |
| Corynebacterium afermentans |
| Corynebacterium argentoratense |
| Corynebacterium callunae |
| Corynebacterium casei |
| Corynebacterium ihumii |
| Corynebacterium jeikeium |
| Corynebacterium matruchotii |
| Corynebacterium mycetoides |
| Corynebacterium pseudodiphtheriticum |
| Corynebacterium pseudogenitalium |
| Corynebacterium resistens |
| Corynebacterium singulare |
| Corynebacterium tuberculostearicum |
| Corynebacterium ureicelerivorans |
| Corynebacterium xerosis |
| Cutibacterium modestum |
| Cutibacterium namnetense |
| Dolosigranulum pigrum |
| Fannyhessea vaginae |
| Gemella sanguinis |
| Granulicatella adiacens |
| Haemophilus haemolyticus |
| Haemophilus sputorum |
| Hoylesella enoeca |
| Hoylesella nanceiensis |
| Hoylesella timonensis |
| Kingella denitrificans |
| Lachnoanaerobaculum saburreum |
| Lactiplantibacillus plantarum |
| Lactobacillus crispatus |
| Lactobacillus iners |
| Lactococcus lactis |
| Lancefieldella parvula |
| Leptotrichia buccalis |
| Leptotrichia hofstadii |
| Leptotrichia shahii |

| **Commensal species** |
| --- |
| Leptotrichia trevisanii |
| Leptotrichia wadei |
| Massilia putida |
| Massilia timonae |
| Metamycoplasma orale |
| Moraxella osloensis |
| Neisseria elongata |
| Neisseria flavescens |
| Neisseria mucosa |
| Neisseria perflava |
| Oribacterium sinus |
| Phocaeicola salanitronis |
| Porphyromonas endodontalis |
| Porphyromonas gingivalis |
| Porphyromonas somerae |
| Prevotella corporis |
| Prevotella denticola |
| Prevotella intermedia |
| Prevotella melaninogenica |
| Prevotella multiformis |
| Prevotella pallens |
| Prevotella scopos |
| Prevotella veroralis |
| Rothia aeria |
| Rothia dentocariosa |
| Rothia mucilaginosa |
| Schaalia odontolytica |
| Segatella baroniae |
| Segatella oris |
| Segatella salivae |
| Solobacterium moorei |
| Streptococcus ilei |
| Treponema denticola |
| Veillonella atypica |
| Veillonella dispar |
| Veillonella parvula |

Note: The reference-genome sequences were downloaded from NCBI. The recovered sequence-accession inventory (commensal_sequence_accessions.csv) is available at https://github.com/Miaaron/AMR-Respiratory-Metagenomic-Simulation. Sequence accessions are not necessarily assembly accession numbers. The supporting literature was not systematically mapped to every species; the two candidate species without available reference genomes could not be identified from the supplied accession inventory.
